# The TLR7/8 agonist INI-4001 enhances the immunogenicity of a Powassan virus-like-particle vaccine

**DOI:** 10.1101/2024.11.28.625832

**Authors:** Michael W. Crawford, Walid M. Abdelwahab, Karthik Siram, Christopher J. Parkins, Henry F. Harrison, Samantha R. Osman, Dillon Schweitzer, Jay T. Evans, David J. Burkhart, Amelia K. Pinto, James D. Brien, Jessica L. Smith, Alec J. Hirsch

## Abstract

Powassan virus (POWV) is a pathogenic tick-borne flavivirus that causes fatal neuroinvasive disease in humans. There are currently no approved therapies or vaccines for POWV infection. Here, we develop a POW virus-like-particle (POW-VLP) based vaccine adjuvanted with the novel synthetic Toll-like receptor 7/8 agonist INI-4001. We demonstrate that INI-4001 outperforms both alum and the Toll-like receptor 4 agonist INI-2002 in enhancing the immunogenicity of a dose-sparing POW-VLP vaccine in mice. INI-4001 increases the magnitude and breadth of the antibody response as measured by whole-virus ELISA, induces neutralizing antibodies measured by FRNT, reduces viral burden in the brain of infected mice measured by RT qPCR, and confers 100% protection from lethal challenge with both lineages of POWV. We show that the antibody response induced by INI-4001 is more durable than standard alum, and 80% of mice remain protected from lethal challenge 9-months post-vaccination. Lastly, we show that the protection elicited by INI-4001 adjuvanted POW-VLP vaccine is unaffected by either CD4^+^ or CD8^+^ T cell depletion and can be passively transferred to unvaccinated mice indicating that protection is mediated through humoral immunity. This study highlights the utility of novel synthetic adjuvants in VLP-based vaccines.

**Author summary:** Powassan virus (POWV) is an emerging pathogenic tick-borne flavivirus for which there is no vaccine. Current tick-borne flavivirus vaccines are less than ideal and use formalin-inactivated virus adjuvanted with alum. These vaccines require thorough inactivation of the antigen and frequent boosting to maintain immunity. In this study, we describe the development of a POWV vaccine using Powassan virus-like-particles (POW-VLPs) adjuvanted with either of two novel Toll-like receptor (TLR) agonists, the TLR4 agonist INI-2002 or the TLR7/8 agonist INI-4001. We show that INI-4001 enhances the antibody response, reduces POWV neuroinvasion, and elicits full protection from lethal POWV infection in mice prime-boost vaccinated with low doses of POW-VLP. We further show that this protection is mediated by a humoral immune response which is both broader and more durable than a POW-VLP vaccine formulated with alum. These findings demonstrate the effectiveness of the novel synthetic TLR7/8 agonist INI-4001 as an adjuvant for low-dose VLP-based vaccines and the ability of this vaccine platform to improve upon current tick-borne flavivirus vaccine methodology.

## Introduction

Powassan virus (POWV) is a tick-borne flavivirus comprising two lineages of pathogenic virus found in North America and Far Eastern Russia maintained in nature by enzootic cycles between various mammalian hosts and a few key species of ixodid ticks [1–10]. Human infection from tick bites can result in life-threatening neuroinvasive disease with a case fatality rate of ∼12% and long-term neurological sequelae in 50% of survivors [11, 12]. Though cases of human infection have been historically rare, the incidence of POWV cases has been increasing and there is evidence that the virus is emerging in North America [1, 3, 13]. Most cases occur in the northeast and Great Lakes regions of the United States where the two main ixodid tick vectors *Ix. cookei* and *Ix. scapularis* are most abundant [1, 14–16]. However, the geographical distribution of POWV cases may expand as the climate continues to warm and POWV vectors continue to spread [14, 16–18]. There are currently no approved vaccines or therapies for POWV infections. These factors make POWV a growing public health concern for which the development of a vaccine should be prioritized.

Virus-like particle (VLP)-based vaccines are a promising platform for the development of a POWV vaccine. VLPs are recombinant vaccines of viral structural proteins that self-assemble into particles that resemble the virus itself, including critical quaternary epitopes ([19, 20] and reviewed in [21–24]). Because they are not capable of replication, VLP-based vaccines are safer than inactivated or attenuated viruses which can pose a health risk to vaccinees if not completely inactivated or attenuated. Flavi-VLPs can be generated through the expression of the viral structural proteins pre-membrane (prM) and envelope (E), the latter being the main target of neutralizing antibody responses to flaviviruses. The resulting presentation of E in a near-native geometry and conformation can surpass formaldehyde-inactivated virus in its ability to induce neutralizing antibodies to structural epitopes [25, 26]. VLPs have been shown to induce not only robust, high-quality humoral responses but also cellular immunity characterized by a Th1 and cytotoxic T lymphocyte (CTL) response making them ideal antigens for future tick-borne flavivirus-vaccines ([21, 27–29] and reviewed in [30, 31]).

Adjuvants are important in generating an effective and durable response to VLPs by enhancing processes that lead to an improved adaptive immune response [32, 33]. Adjuvants accomplish this by altering the uptake of antigens and/or triggering an innate immune response in the absence of infection. Alum is the most common vaccine adjuvant and has a number of proposed methods of action including activating the pattern recognition receptors (PRRs) cGAS-STING and NLRP3 as well as forming a gel-like structure that retains the antigen at the injection site to promote a sustained immune response, though these methods of action are debated (reviewed in [34]). While alum has dominated the field for decades, there are many other PRR-targeting adjuvants that have shown great potential in preclinical and clinical vaccine development for their ability to elicit robust, durable antibody responses as well as T-helper type 1 (Th1) responses. Among these are Toll-like receptor (TLR) agonists which mimic the pathogen-associated molecular patterns recognized by TLRs, a key class of PRR that mediates the initial innate immune and subsequent adaptive immune response to infection. TLR agonists have emerged as particularly effective in the development of virus vaccines. The TLR7/8 agonist INI-4001 (TLR7 agonist in mice and TLR7/8 agonist in humans [35, 36]) and the TLR4 agonist INI-2002 are two novel synthetic adjuvants that have proven effective at improving the humoral and cellular immune response in multiple candidate vaccines making them great candidates for the development of a POWV vaccine [35, 37–40].

In this study, we describe the immunogenicity of a VLP-based vaccine for POWV adjuvanted with either alum, INI-2002, or INI-4001. We demonstrate that the addition of the TLR7/8 agonist INI-4001 significantly improves the neutralizing antibody response and protection against lethal infection from both lineages I and II of POWV. Vaccine adjuvanted with INI-4001 also significantly reduces the viral burden in the brain of infected mice and increases the durability of humoral immunity which appears to mediate protection against challenge as demonstrated by passive transfer. We conclude that POW-VLP adjuvanted with INI-4001 is a highly promising vaccine candidate against POWV.

## Results

### A low-dose POW-VLP vaccine adjuvanted with the TLR7/8 agonist INI-4001 induces a neutralizing antibody response and protects mice from lethal POWV challenge

Previously, we produced a cell line that expresses lineage 1 POWV LB strain (POWV-I) prM-E when induced with doxycycline [41]. Expression of POWV-prM-E leads to the self-assembly and secretion of Powassan virus-like particles (POW-VLP) into tissue culture supernatant, which can be collected and purified. We confirmed the secretion of particles with an average diameter of 25nm by electron microscopy (Fig 1A) and quantified these particles relative to infectious POWV by western blot of E protein (Fig 1B) [42]. We previously demonstrated that prime-boost vaccination of mice with 2X10^7^ E-protein focus-forming-unit equivalents (FFUe) of POW-VLP adjuvanted with alum induced neutralizing antibodies in mice and conferred 100% protection from lethal challenge with POWV-I [41].

**Fig 1.**
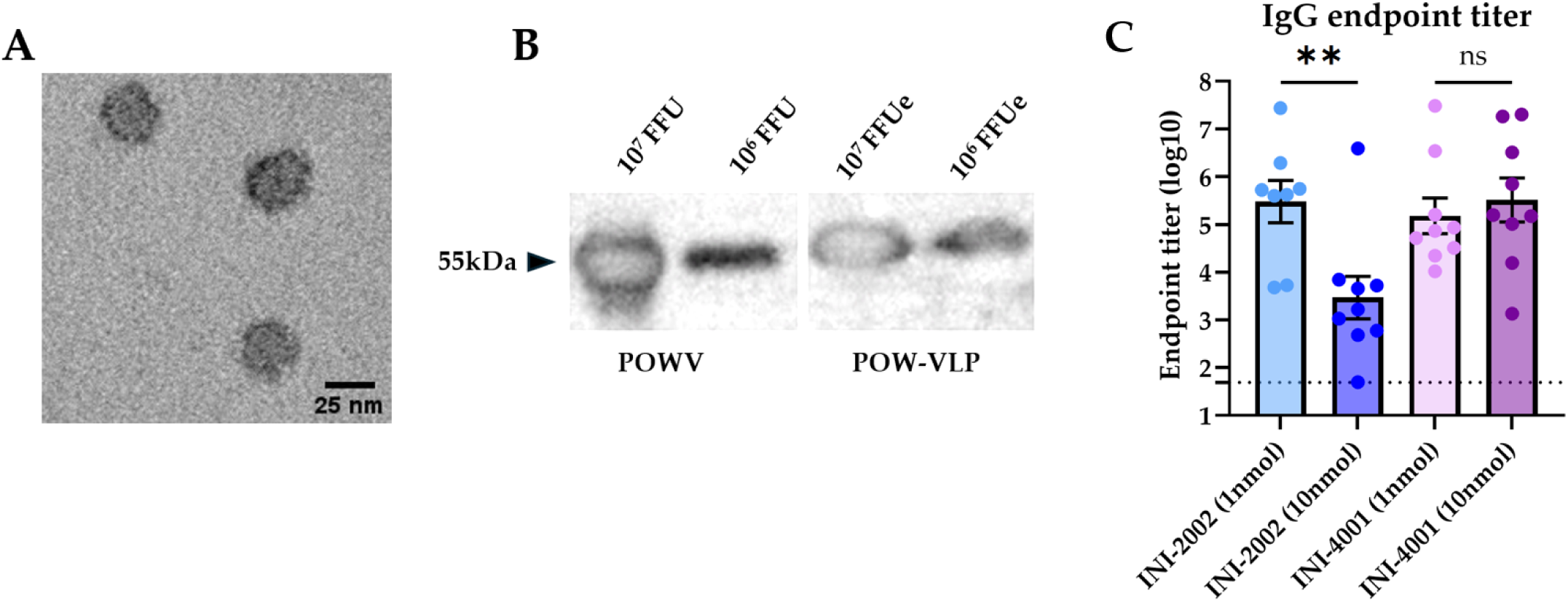
Quantifying VLPs and optimizing adjuvant concentrations for vaccination. (A) Purified POW-VLP visualized by transmission electron microscopy. (B) SDS-PAGE and western blot of POWV (left) and POW-VLP (right) with T077 anti-TBEV-E antibody. Expected migration at 54 kDa. (C) Mice prime-boost vaccinated 2 weeks apart *sc* with 10^6^ FFUe of VLP adjuvanted with either varying concentrations of INI-2002 or INI-4001, n=8-9/group. Serum collected 2 weeks post-vaccination and analyzed by whole-virus ELISA with anti-mouse IgG. Endpoint titers are log-transformed and reported as means ± SEM. Dotted line represents the least dilute serum tested (1:50). Statistical significance determined by one-way ANOVA and Šídák’s multiple comparisons test to compare doses within adjuvant treatment groups. Data represent two independent experiments.

We sought to test whether the TLR4 agonist INI-2002 or the TLR7/8 agonist INI-4001 could improve our POW-VLP vaccine and allow us to reduce the antigen dosage while maintaining immunogenicity and protection. Reduction of antigen also provides greater sensitivity to evaluating adjuvant efficacy. To first determine the appropriate amount of adjuvant for vaccination, we vaccinated C57BL/6 (B6) mice twice subcutaneously (*sc*) two weeks apart with 1X10^6^ FFUe of VLP (20-fold less antigen than we had previously used with alum) adjuvanted with either 1 or 10nmol of either INI-2002 or INI-4001 in their aqueous formulations. We then collected sera 2 weeks post-boost and measured endpoint IgG titer by whole-virus binding ELISA. INI-2002 generated a higher anti-POWV IgG antibody titer at the lower dose of 1nmol while INI-4001 induced equal IgG responses at both doses (Fig 1C). Previous studies with INI-4001 have shown that formulations with higher dosages improve the immune response [35, 38]. Given this information, we chose 1nmol INI-2002 and 10nmol INI-4001 as the working concentrations for subsequent POWV vaccine studies.

Next, we prime-boost vaccinated mice in the same manner as above with either VLP alone or VLP adjuvanted with alum, INI-2002, or INI-4001. Control mice were vaccinated with PBS vehicle alone. Sera collected 2 weeks post-boost vaccination were analyzed for whole-virus binding by ELISA. The mean log IgG endpoint titer in alum-vaccinated mice (3.1) was not significantly different than the background titer observed in mock-vaccinated animals (2.3) or in mice vaccinated with VLP alone (2.5) (Fig 2A). Between the two novel TLR agonists, only INI-4001 produced significantly higher IgG titers (4.9, p = 0.014) than alum. We then measured the neutralizing antibody titers of vaccinated mice and found that only 2/18 of the mice vaccinated with INI-2002 produced focus reduction neutralization test 50 (FRNT50) above the limit of dilution compared to 11/18 of the INI-4001-vaccinated mice (Fig 2B). Together these data demonstrate that INI-4001, but not INI-2002, outperforms alum in inducing IgG and neutralizing antibodies targeting POWV in a low-dose prime-boost POW-VLP vaccine.

**Fig 2.**
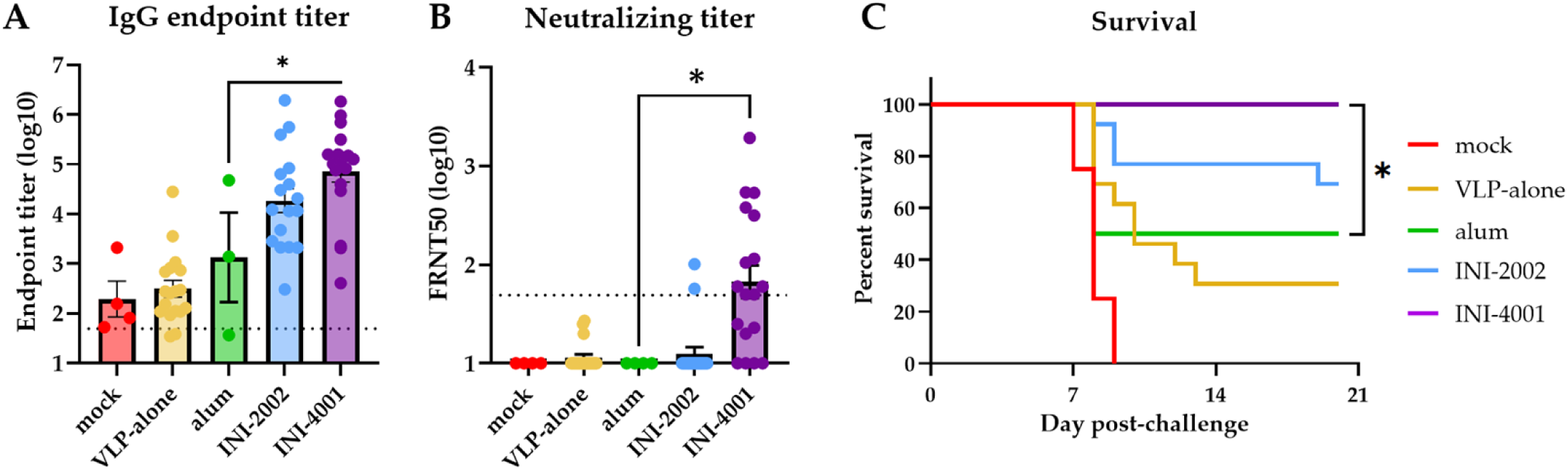
INI-4001-adjuvanted VLP elicits superior antibody response and protection from lethal POWV challenge. (A-B) Mice prime-boost vaccinated 2 weeks apart *sc* with 10^6^ FFUe of VLP alone or adjuvanted with 300µg alum, 1nmol INI-2002, or 10nmol INI-4001. Mock group injected with PBS vehicle alone. mock n=4; VLP-alone n=18-19; alum n=3-4; INI-2002 n=18; INI-4001 n=18. Serum collected 2 weeks post-boost and analyzed by whole-virus ELISA (A) or FRNT (B). Data are reported as log-transformed mean endpoint titer ± SEM for IgG titer and reciprocal FRNT50 for neutralizing titer. Dotted line represents least dilute sera tested (1:50). Data represent three independent experiments. (A) Statistical significance determined by one-way ANOVA and Šídák’s multiple comparisons test to compare treatments to alum group after log transformation. * = p < 0.05. (B) Statistical significance determined by one-way ANOVA and Šídák’s multiple comparisons test to compare alum group to INI-2002 and INI-4001. (C) Vaccinated mice challenged with a lethal 10^4^ FFU dose of POWV-LB 2 weeks post-boost. Survival of mice assessed to day 20 post-challenge. mock n=4; VLP-alone n=13; alum n=4; INI-2002 n=13; INI-4001 n=13. Data represent three independent experiments. Statistical significance determined by log rank test with Bonferroni correction to compare adjuvant groups to VLP-alone. * = p < 0.05.

To determine whether these formulations could elicit protection from POWV, we challenged mice intraperitoneally (*ip*) at the time of serum collection with a lethal dose (10^4^ FFU) of POWV-I. We monitored mice for weight loss and other signs of disease for three weeks. Mice that exhibited clinical signs of morbidity including paralysis and/or substantial weight loss (≥ 20% original body weight) were humanely euthanized. All mock-vaccinated mice died or were euthanized 7-9 days post-infection consistent with previous studies (Fig 2C) [41, 43–46]. Survival of mice that received VLP alone with no adjuvant was 31%, whereas mice that received alum adjuvanted vaccine had a survival rate of 50%. Survival of INI-2002 vaccine mice was higher at 69% but this was not significantly different than the survival of alum-vaccinated mice. Mice vaccinated with INI-4001-adjuvanted POW-VLP were completely protected over the course of the study (100% survival, p = 0.025 compared to alum). Taken together, these data demonstrate that POW-VLP adjuvanted with the TLR7/8 agonist INI-4001 induces both higher IgG and neutralizing antibody titers than a standard alum formulation and confers significant improvement in protection from lethal challenge at this early timepoint.

### INI-4001 decreases viral burden in brain, liver, and spleen of infected mice

POWV is neurotropic and can be detected in the brain of infected humans and mice [44–49]. Similar to infection with the closely related tick-borne encephalitis viruses (TBEV), the neuropathologies associated with POWV are thought to be mediated both by viral neuroinvasion and cytotoxic CD8+ T cell infiltration into the central nervous system [41, 50]. It is therefore important for a vaccine to reduce POWV neuroinvasion. To determine whether POW-VLP adjuvanted with either INI-2002 or INI-4001 reduces dissemination of virus to the brain and other tissues, we infected vaccinated mice with a lethal dose of POWV-I and harvested the brain, liver, spleen, and blood at 6 days post-infection. We then measured viral RNA titer by RT-qPCR in homogenized tissues. Four of the five mock-vaccinated mice had substantial levels of POWV viral RNA in the brain at the time of harvest while INI-4001-vaccinated mice appeared to be largely protected from neuroinvasion with a significantly lower mean titer (Fig 3). Although INI-2002-vaccinated mice trended towards lower POWV burden in the brain, this was not statistically significant. Similarly, INI-4001-vaccinated mice had significantly reduced viral loads in the liver and spleen compared to mock-vaccinated mice with no significant differences detected in the INI-2002-vaccinated mice. These results demonstrate that adjuvantation with INI-4001 reduces viral loads in multiple tissues post-infection, most prominently in brain tissue, consistent with the observed protection against POWV-induced mortality elicited by this formulation.

**Fig 3.**
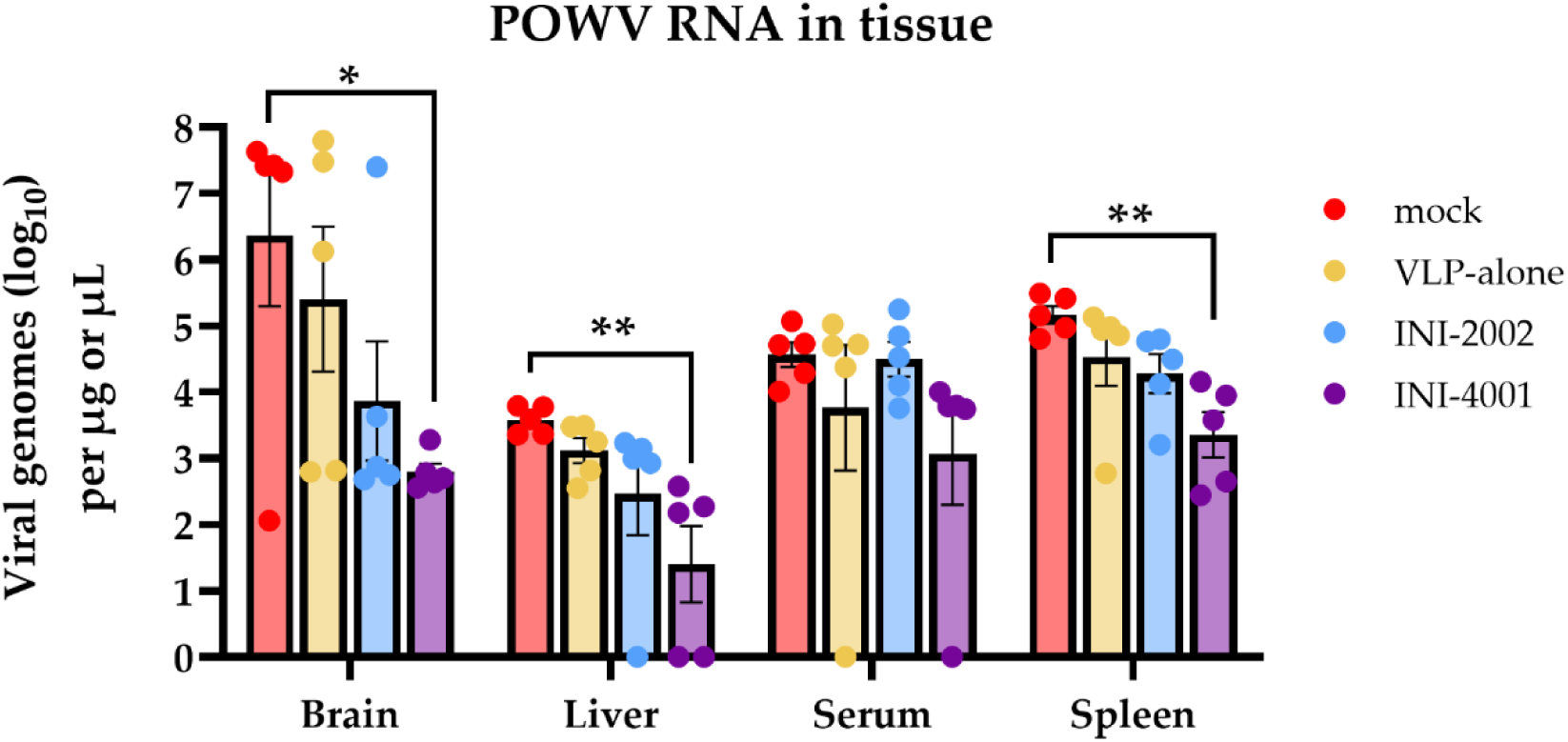
Vaccination with INI-4001-adjuvanted VLP decreases viral RNA burden in multiple tissues post-challenge. Mice prime-boost vaccinated 3 weeks apart with PBS, VLP alone, or adjuvanted VLP. n=5/group. Vaccinated mice infected *ip* with 10^4^ FFU POWV-LB 4 weeks post-boost. Viral load was measured in tissues harvested 6 days post-infection by RT-qPCR. Data are reported as log-transformed mean ± SEM. Statistical significance was determined on log-transformed data using one-way ANOVA with Šídák’s multiple comparisons to mock-vaccinated mice within each compartment.

### Passive transfer of sera from vaccinated mice protects naïve mice from challenge

Given the superior vaccine response and protection elicited by the adjuvant INI-4001 compared to INI-2002, we chose to continue to evaluate our POW-VLP vaccine formulated with INI-4001. We next sought to evaluate how the humoral and cellular arms of the adaptive immune response are contributing to the protection elicited by our vaccine. We first assessed the role of antibody-mediated protection by passively transferring the pooled sera of vaccinated mice into unvaccinated mice one day prior to lethal challenge. IgG and neutralizing antibody titers were measured in pooled sera and in recipient mice the day after transfer (Figs 4A and B). Mice that received sera from vaccinated mice had FRNT50 values very close to the mean measured in INI-4001-vaccinated mice in previous experiments (Figure 2B). Mice that received pooled sera from mock-vaccinated mice died 6-7 days post-challenge while four of the five mice that received vaccinated-mouse sera survived (Fig 4C). Interestingly, the one mouse that succumbed to infection that received vaccinated-mouse serum didn’t begin to decline until day 13 post-challenge (Fig 4D). Given that the half-life of mouse antibodies is on the order of one week, these data suggest that the transferred sera antibodies did not fully clear infection and the mouse succumbed after antibody titers declined below the protective threshold [51]. This experiment demonstrates that the antibody response elicited by our INI-4001-adjuvanted POW-VLP vaccine contributes to protection from lethal challenge.

**Fig 4.**
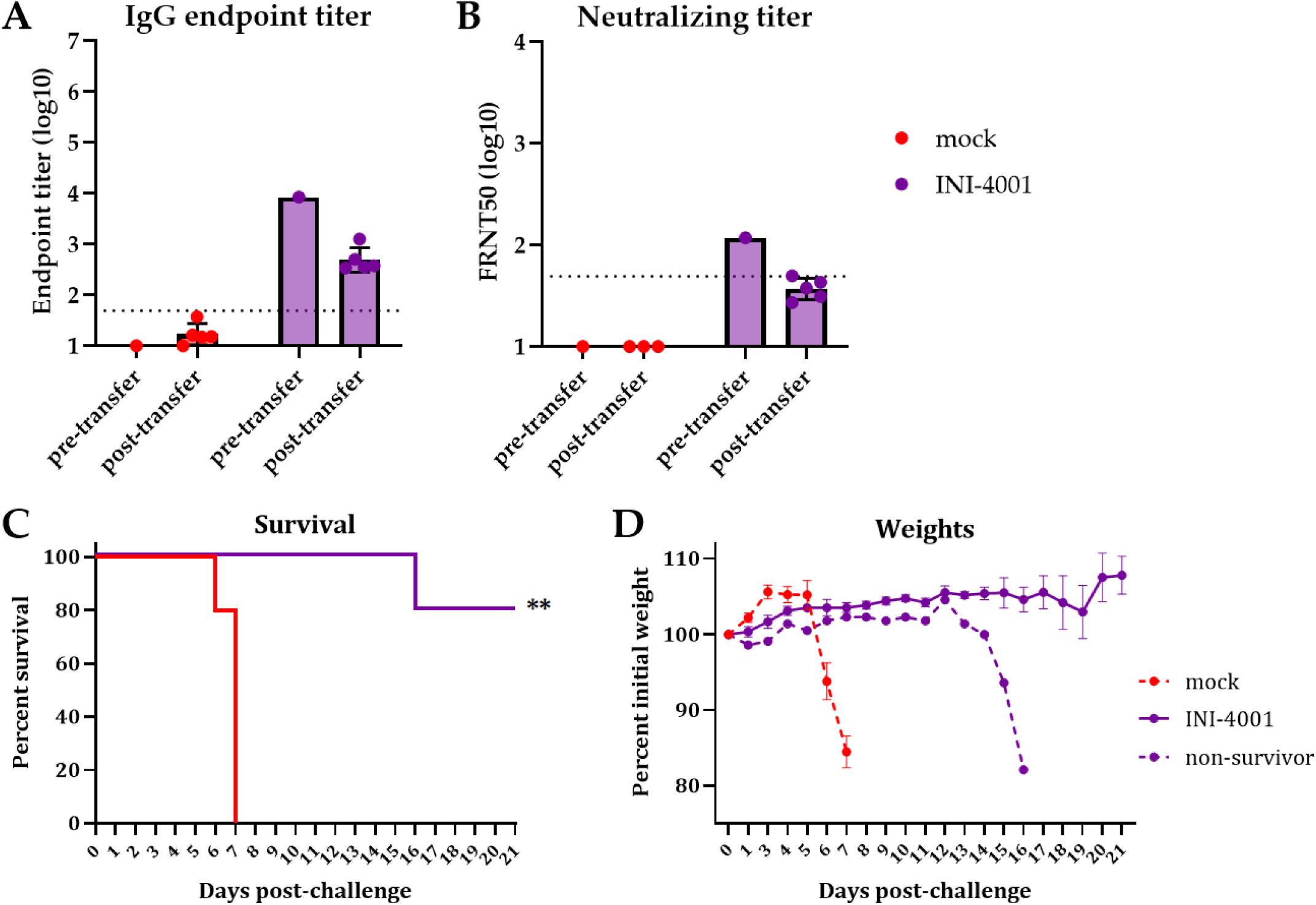
Passive transfer of vaccinated mouse sera protects naïve mice from lethal challenge. Mice vaccinated twice 3 weeks apart. Sera collected 3 weeks post-boost and pooled. n=5/group for both vaccinated donors and unvaccinated recipients. Unvaccinated mice received 200uL of pooled sera *ip* one day before *ip* infection with 10^4^ FFU POWV-LB. Serum collected one day post-transfer from recipient mice and analyzed alongside pre-transfer pooled sera by whole-virus ELISA (A) and FRNT (B). Data are reported as log-transformed mean endpoint titer ± SEM for IgG titer and reciprocal FRNT50 for neutralizing titer. Dotted line represents least dilute sera tested (1:50). Survival (C) and weights (D) monitored for 21 days post-challenge. (C) Statistical significance determined by log rank Mantel-Cox test to compare vaccinated animals to mock. ** = p < 0.01. (D) Weights represented as mean percent of initial weights ± SEM. All data represent single experiment.

### Depletion of CD4+ or CD8+ T cells does not affect protection

The lack of measurable neutralizing titers in several mice that are protected from lethal POWV challenge (Figure 2B & C) raised the question of whether a portion of the vaccine-induced protection is mediated by non-humoral adaptive immunity. Although we were unable to detect POWV-specific T cell responses in vaccinated mice (S1 Fig), we assessed whether T cells are required for mediating protection by performing T cell depletion in vaccinated animals prior to challenge and observing differences in survival. We depleted CD4^+^ or CD8^+^ T cells by administering CD4- or CD8-depleting antibodies in prime-boost vaccinated mice 3 days prior to and the day of challenge (Fig 5A). We confirmed T cell depletion by flow cytometry of blood 3 days post-challenge (Figs 5B-D). Survival of the infected mock-vaccinated animals was not impacted by either CD4^+^ or CD8^+^ T cell depletion compared to the isotype control group (Fig 5E), nor were there differences in weight loss for these animals (Fig 5F). Similarly, neither CD4-nor CD8-depletion affected the survival of INI-4001-vaccinated mice, with 100% of all mice surviving lethal challenge and maintaining weights regardless of depleting antibody. These data suggest that T cells are not necessary for the protection against POWV challenge that is elicited by POW-VLP adjuvanted with the TLR7/8 agonist INI-4001 at this early timepoint.

**Fig 5.**
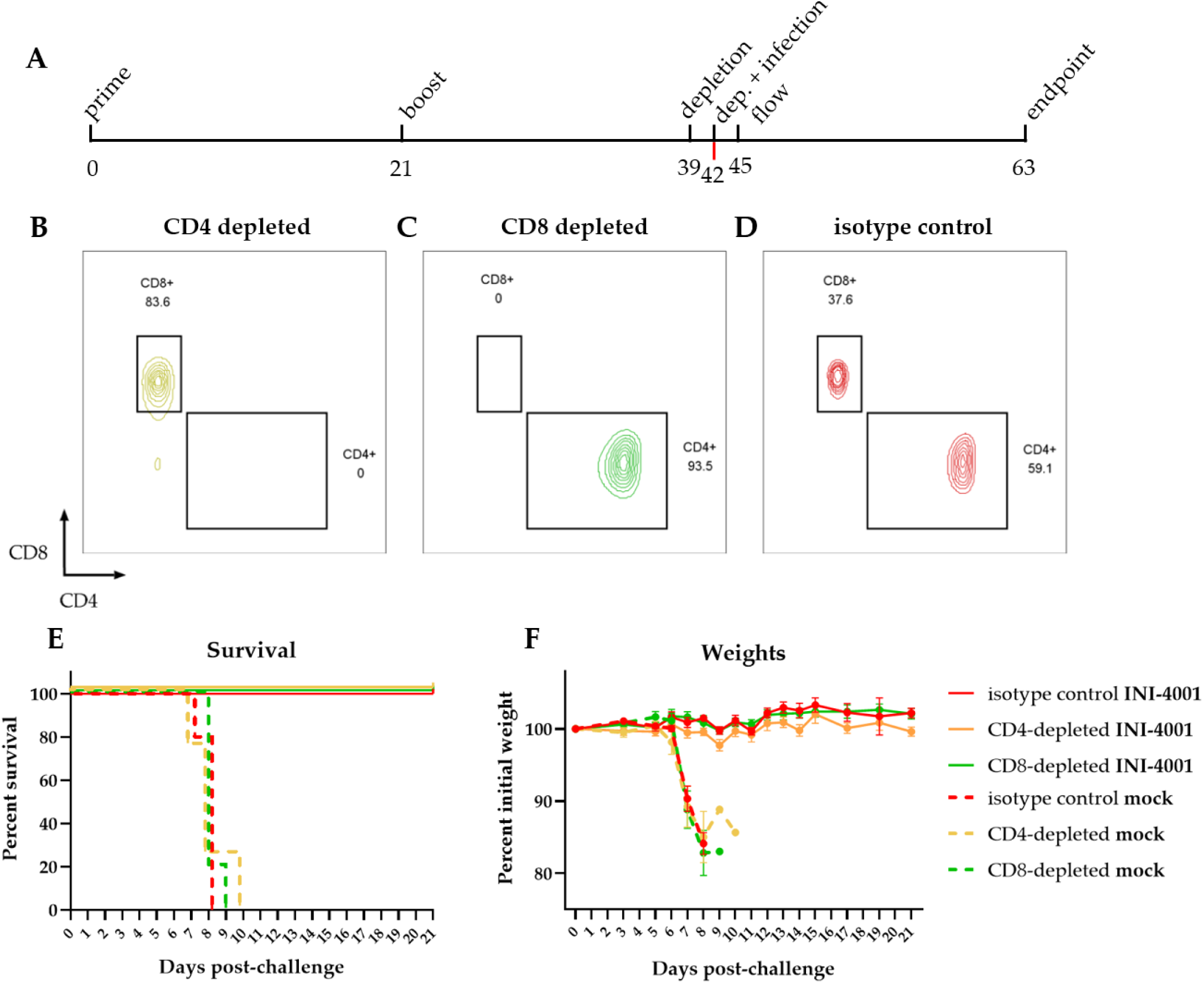
T cell depletion does not affect survival of vaccinated mice. (A) Timeline of experiment in days. Mice were prime-boost vaccinated 3 weeks apart (days 0 and 21) *sc* with 10^6^ FFUe of POW-VLP adjuvanted with 10nmol INI-4001. The mock group was injected with PBS vehicle alone. Mice were then treated *ip* with 100 µg of either CD4-depleting, CD8-depleting, or non-depleting isotype control (n=5/group) 3 weeks post-boost (day 39) and again three days later (day 42). Mice were challenged with a lethal 10^4^ FFU dose of POWV-LB *ip* at time of second depletion (day 42). Blood was collected by tail vein 3 days after second administration of depleting antibodies to confirm T cell depletion (day 45). Survival and weights of mice monitored to day 21 endpoint post-challenge (day 63). (B-D) Representative flow plots of T cells from (B) CD4-, (C) CD8-, or (D) control-depleted mice 3 days post-depletion. Cells gated on CD45^+^ CD3^+^ CD19^−^ CD4^+^/CD8^+^. (E-F) Survival (E) and weights (F) of mock (dashed lines) or INI-4001 (solid lines) vaccinated mice after depletion of either CD4^+^ (yellow) or CD8^+^ (green) T cells compared to isotype control (red). (E) Statistical significance determined by log rank test with Bonferroni correction to compare depleted groups to isotype control. No significant differences were found. (F) Weights represented as mean percent of initial weights ± SEM. Data represent one experiment.

### Vaccine against POWV-I adjuvanted with INI-4001 agonist generates cross-reactive antibodies to other tick-borne flaviviruses and protects against lethal challenge from POWV-II

There are two lineages of POWV, both of which infect and cause disease in humans. POWV-LB is the prototype of the lineage I (POWV-I) whose genome we used to generate our POW-VLP and the virus used for challenge studies thus far. However, protection against both lineages is important to protect individuals against POWV disease. We therefore sought to evaluate the cross-reactivity of antibodies from vaccinated mice to the prototypical POWV-II Spooner strain as well as another more distantly related tick-borne flavivirus, Langat virus (LGTV), and an even more distantly related mosquito-borne flavivirus, West Nile virus (WNV). Vaccination with INI-4001 induced antibodies that could bind to both POWV-II and LGTV by ELISA, while alum did not generate any measurable cross-reactive binding activity (Fig 6A). Neither alum-nor INI-4001-vaccinated mice produced antibodies that could cross-react to WNV.

**Fig 6.**
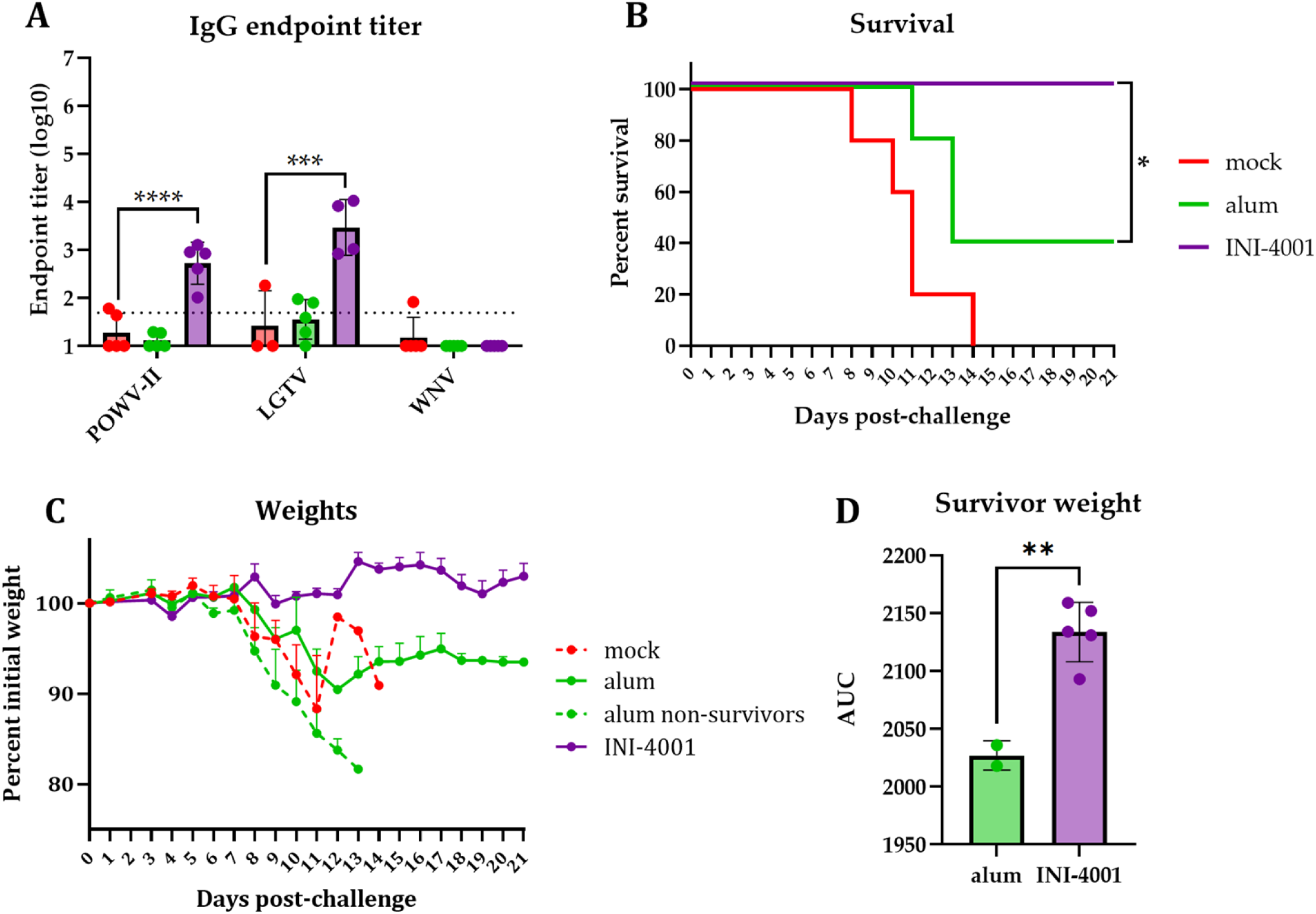
Vaccination with INI-4001-adjuvanted POWV-I-VLP generates antibodies that bind POWV-II and LGTV and protects mice from lethal POWV-II challenge. Mice prime-boost vaccinated 3-4 weeks apart *sc* with 10^6^ FFUe of VLP alone or adjuvanted with 300µg alum or 10nmol INI-4001. n=5/group. Mock group injected with PBS vehicle alone. (A) Serum collected 3 weeks post-boost and analyzed by whole-virus ELISA. Data are reported as log-transformed mean endpoint titer ± SEM. Statistical significance determined by one-way ANOVA with Šídák’s multiple comparisons test to compare vaccinated mice to mock-vaccinated mice. Data for POWV-II and LGTV represent one experiment; data for WNV represent independent experiment. n=5/group except for LGTV n=3 for mock and n=4 for INI-4001. (B-C) Mice challenged *ip* 3 weeks post-boost with 10^5^ FFU dose of POWV-II. Survival (B) and weights (C) monitored for 21 days post-challenge. (B) Statistical significance determined by Mantel-Cox log rank test to compare alum and INI-4001 groups. * = p = 0.05. (C) Weights represented as mean percent of initial weights ± SEM. (D) Area under the curve (AUC) of individual alum and INI-4001 survivors. Statistical significance determined by Student’s unpaired t-test. ** = p < 0.01. Data represent one experiment.

We next sought to evaluate whether POW-VLP adjuvanted with INI-4001 could protect from POWV-II challenge. Mock-vaccinated mice infected with a lethal dose (10^5^ FFU) of POWV-II all succumbed to infection by day 14 post-challenge with a median survival of 11 days (Fig 6B), as observed previously [41]. All five mice that received the INI-4001-adjuvanted vaccine survived POWV-II challenge while only two of five alum-vaccinated mice survived this challenge, which was a statistically significant difference. Both alum-vaccinated survivors lost 10% of their body weight before partially recovering back to ∼93% of their initial weight (Fig 6C). Comparatively, the largest weight drop within the INI-4001-vaccinated group was ∼3%, and the mean ranged from 99-105% and finished at 103%. Area under the curve analysis was performed on the surviving mice that received either alum- or INI-4001-adjuvanted VLP which revealed a statistically significant difference in weights between these groups over the course of the challenge (p = 0.03; Fig 6D). In conclusion, INI-4001 further outperforms alum as a vaccine adjuvant by inducing the production of cross-reactive antibodies to both POWV-II and LGTV and protects mice from lethal POWV-II challenge.

### INI-4001 elicits a durable immune response

We lastly sought to evaluate the durability of the immune response elicited by alum, INI-2002, or INI-4001. To do this, we measured the IgG and neutralizing antibody titers in the sera from vaccinated mice every month for 8 months (36 weeks) post-boost vaccination. Anti-POWV-I IgG antibody titers induced by all three adjuvanted vaccines peaked 8 weeks post-boost (Fig 7A) with peak log endpoint titers that were nearly 1,000-fold higher in mice that received INI-4001-compared to alum-adjuvanted VLP. By 20 weeks post-boost, the IgG titers in all adjuvant groups dropped to a lower plateau that was maintained to the last timepoint. While INI-2002-vaccinated mice had significantly higher titers than alum-vaccinated mice at multiple timepoints, INI-4001-vaccinated mice maintained IgG titers that were significantly higher than alum-vaccinated mice throughout the experiment.

**Fig 7.**
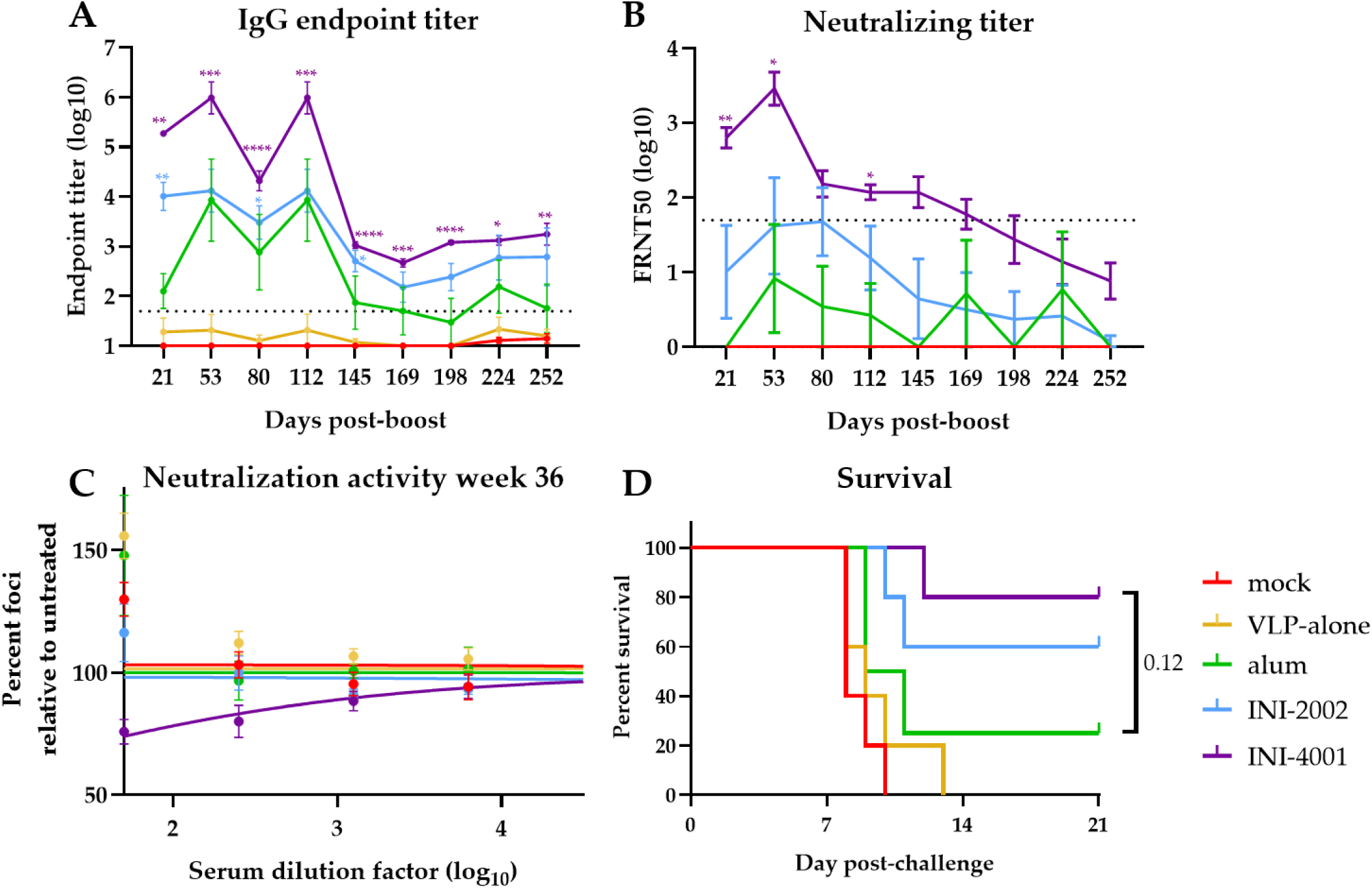
INI-4001-adjuvanted vaccine elicits durable antibody response with long-lasting protection. (A-B) Mice vaccinated twice 4 weeks apart. Serum collected monthly for ∼9 months post-boost and analyzed by whole-virus ELISA for IgG titer or reciprocal FRNT50 assay for neutralizing titers. (A,B) Data are log transformed and reported as means ± SEM. Statistical significance was calculated by repeated measure two-way ANOVA and Šídák’s multiple comparisons test to compare TLR agonist treatments to alum-treated group after log transformation: * = p < 0.05; ** = p < 0.01; *** = p < 0.001; **** = p < 0.0001. Dotted line represents least dilute sera tested (1:50). (C) Neutralization activity of sera 36 weeks post-boost as measured by percent of foci per well of serum-treated virus relative to untreated control. Data represent treatment-group mean percentages ± SEM. Lines represent non-linear regression curves. (D) Mice infected *ip* at 9 months post-boost with 10^4^ FFU POWV-LB. Survival monitored for 21 days. Statistical significance determined by log rank test with Bonferroni correction to compare adjuvant groups to alum treatment. All data represent single experiment. n=4 for alum; n=5/group for all other groups.

The neutralizing antibody responses in adjuvant-vaccinated animals also peaked by 8 weeks post-boost before slowly declining in all groups (Fig 7B). FRNT50 values peaked at dilutions of 1:8 for alum-vaccinated mice, 1:48 for INI-2002, and 1:2,890 for INI-4001. By the end of the experiment, none of the alum-vaccinated mice had measurable neutralizing antibodies, and only one of the five INI-2002-vaccinated mice had a measurable titer. In contrast, four of the five INI-4001 mice had measurable titers. Though these final FRNT50 values fell below the limit of dilution and are therefore estimates, neutralization was readily observable in the sera of the INI-4001 group (Fig 7C). Throughout the experiment, INI-4001-adjuvanted VLP was the only vaccine that elicited mean neutralizing titers significantly higher than alum at any timepoint.

To test if these vaccine formulations induced durable protection, mice were challenged with POWV at 36 weeks post-boost. Only one of the four alum-vaccinated mice survived challenge compared to three of the five INI-2002-vaccinated mice and four of the five INI-4001-vaccinated mice (Fig 7D). Although the probability of survival in the INI-4001 vaccine group was not significantly different than in the alum group (p=0.095), this may be due to the small number of mice used in this study (n=4-5). Taken together, these results suggest that although the diminishing kinetics of the antibody responses were similar between vaccines, the higher initial response elicited by INI-4001-adjuvanted VLP creates a more durable antibody response and this translates to long-lasting protection from challenge nearly 9 months post-vaccination.

## Discussion

POWV is a significant growing public health concern in North America for which there are currently no medical interventions. With an infection case fatality rate of 12% and a dramatic increase in the number of cases over the past two decades, the development of an effective and durable vaccine against POWV is critical [11, 12, 52]. There are several effective vaccines for other closely related tick-borne flaviviruses (TBFVs) including six approved vaccines for the tick-borne encephalitis viruses (TBEVs) and one for Kyasanur Forest disease virus [19–21]. These vaccines are all formalin-inactivated viruses, and those for TBEV are adjuvanted with alum. Though these vaccines are well-tolerated, immunogenic, and efficacious (reviewed in [19, 22, 23]), they have shortcomings including the risk posed by incomplete inactivation of virus used in vaccines, the possibility that important epitopes might be modified during inactivation, and the requirement for boosting every 1-5 years to maintain protective neutralizing titers [19–21, 24–26]. These points highlight the need to improve our current methodology for TBFV vaccines.

VLP-based vaccines have emerged as safe alternative vaccine candidates that maintain neutralizing epitopes and are thus expected to induce potent immune responses. Additionally, novel adjuvants such as TLR agonists have emerged as useful tools for improving vaccine responses and durability (reviewed in [53]). Here, we demonstrate the potent combination of POW-VLP and the novel TLR7/8 agonist INI-4001 in the design of a POWV vaccine in mice. Prime-boost vaccination using a low-dose of 10^6^ FFUe of POW-VLP adjuvanted with INI-4001 rapidly induced a significantly higher anti-POWV IgG and neutralizing antibody titer in mice compared to VLP adjuvanted with a standard alum formulation (Figs 2A-B) and conferred 100% protection from lethal POWV challenge (Fig 2C) validating the utility of this novel adjuvant.

The neuropathology associated with encephalitic TBFVs such as POWV is believed to arise from central nervous system (CNS) inflammation and leukocyte invasion following viral replication in the brain and CNS [44]. Mice that received INI-4001-adjuvanted vaccine had significantly reduced viral RNA loads in the brain, consistent with the protection from lethal challenge and absence of neurological symptoms in these mice (Fig 3). INI-4001-vaccinated mice also had reduced viral RNA loads in the liver and spleen. Interestingly though, there was no difference in serum viremia between vaccinated and unvaccinated animals suggesting that tissue burden and viremia are independent at this timepoint. It is possible that earlier differences in viremia may have influenced the subsequent viral burden in these tissues, but that these differences were no longer apparent 6 days post-challenge. Santos *et. al* and Reynolds *et. al* have shown that POWV viremia in mice peaks much earlier at about 1-2 days post-infection which supports this possibility [44, 46].

The need to develop TBFV vaccines that require less frequent boosting to maintain protection led us to test whether the vaccine response elicited by INI-4001-adjuvanted VLP was more durable than that of VLP adjuvanted with alum. Indeed, we found that POW-VLP adjuvanted with INI-4001 induced a more durable IgG and neutralizing antibody response (Fig 7A-B). Furthermore, vaccination with INI-4001-adjuvanted VLP protected 80% of mice from lethal infection 36 weeks post-boost compared to only 25% with alum-adjuvanted VLP (Fig 7C). The enhanced durability of the vaccine response induced by INI-4001 demonstrates the potential to improve our approach for both current and future TBFV vaccines.

The presence of two similar but genotypically distinct lineages of POWV, lineages I and II (POWV-I and POWV-II or deer tick virus) led us to investigate how broad the response to our POWV-I-based VLP vaccine is. We found that mice that received INI-4001-adjuvanted vaccine generated antibodies that bound not only POWV-II which shares 96% amino acid identity in the E protein to POWV-I, but also to LGTV which shares ∼77% identity (Fig 6A). Mice that received alum-adjuvanted vaccines did not generate cross-binding antibodies to either of POWV-II or LGTV demonstrating that INI-4001 uniquely improves the breadth of the antibody response. This improved breadth of the vaccine response translated to 100% protection from lethal POWV-II challenge in mice that received INI-4001-adjuvanted VLP compared to only 40% in those that received alum-adjuvanted VLP (Fig 6B).

Cell-mediated immunity characterized by antigen-specific cytotoxic CD8+ T cells and a Th1 response is ideal for the clearance of intracellular pathogens such as viruses (reviewed in [56]). Enhancing the T cell response to virus vaccines would likely improve efficacy and durability [54, 55]. Previous studies have shown that INI-4001 skews vaccine responses towards Th1 as do other TLR7/8 agonists [35, 38, 56–58]. We therefore sought to determine whether the protection conferred by our vaccine was mediated by humoral or cellular immunity. We found that while passive transfer of the pooled sera of vaccinated mice protected naïve mice from lethal challenge (Fig 4C), neither depletion of CD8^+^ nor CD4^+^ T cells affected vaccine-mediated protection (Fig 5E). We conclude that the protection elicited by our vaccine is therefore antibody mediated at this early timepoint 3-weeks post-boost. Whether the durability of our vaccine is cell mediated is worth future investigation.

Though neutralizing antibodies are considered the correlate of protection for several flaviviruses including the closely related TBEVs [59, 60], it is notable that a measurable neutralizing antibody titer at the time of challenge was not a prerequisite for survival in our study (Figs 2 and 7). There were consistently higher percentages of survivors within each vaccine group than there were animals with measurable neutralizing titers at the time of infection: 31% survival vs 16% measurable FRNT50 for VLP-alone, 50% vs 0% for alum, 69% vs 11% for INI-2002, and 100% vs 78% for INI-4001. There are several possible explanations for this which are not mutually exclusive. First, the neutralizing titer necessary for protection may be lower than the level of detection for our assay. Second, a memory response to infection may raise neutralizing antibody titers to a protective level not seen at the time of serum collection. Work by Uhrlaub *et. al* supports this explanation by demonstrating that the protection elicited by a vaccine against WNV in older mice with minimal neutralizing antibody responses relies on a memory B-cell response to infection [61]. Third, there may be other non-neutralizing mechanisms to the observed protection, either humoral or cellular. Though we did not see a difference in protection after T cell depletion, which would suggest that cell-mediated mechanisms are not responsible for this, all of the INI-4001-vaccinated mice in the T cell depletion experiment had measurable neutralizing titers at the time of infection, so we cannot rule out the possibility that mice with no measurable neutralizing antibodies in previous experiments were protected through cell-mediated immunity. Further investigation into the correlates of protection for POWV are needed.

In conclusion, our study highlights the utility of novel vaccine adjuvants in improving the protective immunogenicity of a VLP-based vaccine targeting POWV. We demonstrate that the TLR7/8 agonist INI-4001 outperforms alum as an adjuvant in multiple facets of the vaccine response elicited by a dose-sparing POWV-VLP vaccine. Further studies investigating whether combinations of INI-4001 alongside other adjuvants including alum and TLR4 agonists can improve upon this design are warranted. Further characterization of INI-4001 as well as other nanoparticle-based vaccine formulations is underway to better understand how this novel adjuvant induces such a potent protective immune response. Additional future experiments will focus on using this infection model and vaccine platform to interrogate the correlates of protection for POWV disease and how best to induce these responses for the development of a vaccine.

## Methods

### Mouse experiments

5-week old male C57BL/6 mice were purchased from Jackson Laboratory and housed in an ABSL-3 facility at Oregon Health & Science University’s (OHSU) Vaccine and Gene Therapy Institute accredited by the Association for Accreditation and Assessment of Laboratory Animal Care (AALAC) in compliance with protocols approved by the OHSU’s Institutional Animal Care and Use Committee (IACUC) #1432.

Primary vaccinations were administered at 7-weeks of age subcutaneously (*sc*) in the dorsal region using 10^6^ FFU equivalents of POW-VLP and either 300µg of Alhydrogel® 2% adjuvant, 1-10nmol of INI-2002, or 1-10nmol of INI-4001 diluted to a final volume of 100uL in sterile injection-grade PBS. INI-2002 and INI-4001 were synthesized as described by Miller *et. al* [35]. An aqueous formulation of INI-4001 was prepared using high-shear homogenization. INI-4001 was weighed into a glass vial, and an adequate volume of an aqueous buffered vehicle containing 50 mM TRIS buffer and 0.1% Tween 80 was added. The mixture was homogenized with a high-shear homogenizer (Silverson L5MA) at 10,000 rpm for 20 minutes. Meanwhile, the aqueous formulation of INI-2002 was prepared by solubilizing it in 2% glycerol using a bath sonicator (FB11201, Fisherbrand, Thermo Fisher Scientific) for 3 hours at a temperature below 35°C. Both INI-4001 and INI-2002 formulations were sterile-filtered using a 13 mm Millex GV PVDF filter with a pore size of 0.22 µm (MilliporeSigma). The formulations were characterized using a Zetasizer Nano-ZS (Malvern Panalytical, Malvern, UK) and exhibited a hydrodynamic particle size of less than 120 nm. The concentrations of INI-4001 and INI-2002 were determined by RP-HPLC according to a previously published method [39]. Mock-vaccinated mice received 100uL of PBS alone. Boost-vaccinations were administered 2- to 4-weeks post-primary vaccination in the same manner. Blood was collected by tail-vein at specified timepoints post-vaccination and left to clot at room-temperature for 30 minutes before centrifugation twice at 10,600 x *g* to collect the serum. Serum was then stored at −20°C until use.

For lethal challenge infections, 10^4^ FFU of POWV LB or 10^5^ FFU POWV Spooner was diluted into 100uL sterile PBS and injected intraperitoneally (*ip*). Mice were monitored for up to three weeks for signs of morbidity including piloerection, hunched posture, ataxia, malaise, paralysis, and weight loss. Moribund mice were defined as those that experienced substantial weight loss ≥ 20% original body weight, paralysis, and/or other signs of morbidity. Moribund mice were euthanized by isoflurane followed by cervical dislocation to limit suffering.

### Reverse transcription-quantitative polymerase chain reaction (RT-qPCR)

For tissue harvest, mice were euthanized with isoflurane and tissues were collected in TRIzol®. Tissues were homogenized by bead-beating with SiLiBeads using a Precellys 24 bead beater homogenizer (Bertin Technologies). Samples were clarified by centrifugation at ∼21,000 x *g* and supernatant RNA was isolated by Quick-RNA Viral Kit (Zymo Research #R1034). RNA was quantified by NanoDrop™ 2000c (ThermoFisher). For blood, serum was collected as previously described and combined with DNA/RNA Shield (Zymo Research #R1100) and processed for RNA using the Quick-RNA Viral Kit. RT-qPCR was performed on 200ng of total RNA from tissue per reaction or 4uL of serum RNA samples using POWV specific primers (Invitrogen: forward TGTTCTGCTGTTCCCGTGAGT; reverse GATGCGCAGCATGTCTTCTG), probe (Applied Biosystems: AGCATCCACGCGAGTG), and TaqMan™ RNA-to-CT™ 1-Step Kit (Catalog #4392656) on an AB StepOne™ Real-Time PCR System (ThermoFisher). Dilutions of known quantities of POWV cDNA were used to produce a standard curve for absolute genome quantitation.

### Passive transfer

Mice were vaccinated as previously described with either INI-4001 adjuvanted vaccine or PBS alone. Sera were collected 3 weeks post-boost, pooled within each group, and heat-treated at 55°C for 30 minutes to inactivate complement. 200µL of pooled serum was passively transferred *ip* to unvaccinated mice. Mice were then challenged with 10^4^ FFU of POWV LB *ip* the day after passive transfer and monitored for three weeks for signs of morbidity and survival.

### T cell depletion and flow cytometry

Mice were vaccinated as previously described. At both days 19- and 21-post boost-vaccination, 100 µg of either CD4-depleting antibody (Clone GK1.5 IgG2b Fisher), CD8-depleting antibody (Clone 2.43 IgG2b Fisher), or non-depleting isotype control (Clone LTF-2 Fisher) was administered *ip*. Mice were then challenged with 10^4^ FFU of POWV LB as previously described. To confirm depletion, 100µL of blood was collected by tail vein 3 days post-infection into 500uL of 100mmol EDTA (Invitrogen), washed with FACS buffer (PBS with 5% FBS and 1mM EDTA), and stained with CD45 AlexaFluor 700 (BioLegend rat anti-mouse clone 30-F11 Cat#103218), CD3 BUV395 (BD Horizon rat anti-mouse Clone 17A2 (RUO) Cat#: 569614), CD19 PE (BioLegend rat anti-mouse clone 1D3/CD19 Cat#: 152408), CD4 V450 (BD Horizon rat anti-mouse clone: RM-4-5 Cat#: 560468), and CD8b PerCP/Cy5.5 (BioLegend rat anti-mouse clone YTS156.7.7 Cat#: 126609) for 30 minutes at room temperature. Cells were then washed and fixed using RBC lysis/fixation solution (BioLegend #422401) for 10 minutes, washed, and resuspend in FACS buffer. Cells were then analyzed by flow cytometry using a BD FACSymphony Spectral Cell Analyzer. T cells were gated as CD45+ CD3+ CD19-.

### Peptide restimulation of T cells

7-week-old mice were vaccinated twice 4 weeks apart and then euthanized 4 weeks post-boost. Spleens were processed to a single-cell suspension over a 40µm cell strainer and suspended in RPMI supplemented with 5% FBS and 1% HEPES. 1× 10^6^ cells were plated per well in a round-bottom 96-well plate and stimulated for 6 hours at 37°C, 5% CO2 in the presence of 10µg/mL brefeldin A and either α-CD3 (clone 2C11) as a positive control, PBS, or 10µg/mL of peptide. Previously defined antigen specific epitopes were used to quantify antigen specific T cells with POWV-E_525-535_ to stimulate CD4^+^ T cells and POWV-E_282-291_ to stimulate CD8^+^ T cells [41]. Following peptide restimulation, cells were washed twice with 1X PBS and stained overnight in 1X PBS at 4°C for the following surface antigens: CD4 (clone RM-4-5), CD8α (clone 53-6.7), and CD19 (clone 1D3). Cells were then washed twice with 1X PBS and fixed in 2% paraformaldehyde at 4°C for 10 minutes. Following fixation, cells were permeabilized with 0.5% saponin and stained in this solution for 1 hour at 4°C for intracellular IFN-γ (clone B27). Following intracellular staining, cells were washed with 0.5% saponin twice, followed by one wash with 1X PBS. Cells were resuspended in 200μl 1X PBS and analyzed by flow cytometry using an Attune NxT focusing flow cytometer. For analysis, T cells were gated on lymphocytes, CD19-, and either CD4-/CD8+ or CD4+/CD8-cells. Antigen specific cells were identified as those producing IFN-γ in the presence of the POWV envelope epitope.

### Cells

HEK293T and Vero E6 cells were maintained in Dulbecco’s Modified Eagle Medium (DMEM) (Corning) supplemented with 5% fetal bovine serum (FBS; HyClone™ #3039603), 100U/mL penicillin, 100 µg/mL streptomycin, and 292 µg/mL L-glutamine (Gibco™ #10378016). HEK293T cells were passaged using citrate buffer. Vero cells were passaged using trypsin 0.05% EDTA (Gibco™ #25-300-120).

### VLP production

The generation of HEK293 cells expressing POWV prM-E was described previously [41]. In short, HEK293 cells were transfected with pLVX-Tet-On Advanced (Takara) to express the tetracycline-controlled transactivator. Tet-transactivator expressing HEK293 cells were then transduced with pseudotyped lentivirus containing pLVX-POWVprME packaged using the packaging vector pSPAX2 and vesicular stomatitis virus G protein expression vector pMD2.G. Transduced cells were selected for by using 1 µg/mL puromycin to generate a HEK293 cell line that expresses POWV-prME when induced with doxycycline.

To produce POW-VLP, HEK293-POWV-prME cells were cultured with 1µg/mL doxycycline in DMEM supplemented with 2% FBS, 100U/mL penicillin, 100 µg/mL streptomycin, and 292 µg/mL L-glutamine. Supernatant was collected 4 days post-induction, centrifuged at 1,000 x *g* for 10min, and filtered through a 0.45µm to remove cellular debris. VLPs were then concentrated and purified by ultra-centrifugation through a 20% sorbitol 50mM Tris 1mM MgCl_2_ pH 8.0 buffer at 150,000 x *g* for 2 hours at 4°C. Pellets were then resuspended in 1/100^th^ of the original volume with 10mM Tris 150mM NaCl buffer supplemented with 5% trehalose for protein stability and stored at −80°C until use. For quantification, VLP preparations were boiled at 95°C for 5 minutes in non-reducing sample buffer. Samples were then run on 10% SDS-PAGE gel, transferred to Immobilon™-P PVDF 0.45µm Membrane (MilliporeSigma #IPVH00010), and immunoblotted with T077 (described later) to visualize E using monkey IgG gamma peroxidase-conjugated antibody (Rockland #617-103-012) and Pierce™ ECL Western Blotting Substrate (ThermoFisher #32106). Pictures were taken using a G:Box imager (Syngene).

### Viruses

POWV LB and Spooner strains were generously provided by Michael S. Diamond (Washington University School of Medicine, St. Louis, MO). LGTV TP21 (NR-51658) was obtained from the Biodefense and Emerging Infections Research Resources Repository (BEI Resources). All viruses were propagated on Vero E6 cells (5 days for LGTV; 7 days for POWV and WNV 385-99 [62]). Supernatant was collected, centrifuged at 1,000 x *g* for 10min, and filtered through a 0.45µm to remove cellular debris. Virus was then concentrated and purified by ultra-centrifugation through a 20% sorbitol 50mM Tris 1mM MgCl_2_ pH 8.0 buffer at 150,000 x *g* for 2 hours at 4°C. Pellets were then resuspended in 1/100^th^ of the original volume in DMEM with 0.1% FBS and stored at −80°C until use. Viruses were titered using limited dilution focus forming assay on Vero cell monolayers. Briefly, cells were infected by rocking for 1 hour at 37°C in 5% CO_2_. Cells were then overlaid with a formula consisting of 2 parts DMEM with 5% FBS, 100U/mL penicillin, 100 µg/mL streptomycin, and 292µg/mL L-glutamine and 1 part 1% high / 1% low viscosity carboxymethylcellulose (CMC) diluted in a 60% PBS aqueous solution. Cultures were then aspirated and fixed at 48 hours post-infection with 4% paraformaldehyde for 30 minutes. Staining protocol to visualize foci was then performed as described in “Focus reduction neutralization test (FRNT)” section of methods. Foci were counted using AID Elispot 7.0 (Autoimmun Diagnostika GMBH) and titers were determined from these numbers.

### Enzyme-linked immunosorbent assay (ELISA)

To titer whole virus binding antibodies, Corning™ Costar™ Brand 96-Well EIA/RIA plates (ThermoFisher) were coated with 10^5^ FFU of POWV-I, LGTV, or WNV or 3.2X10^4^ FFU of POWV-II in 100µL PBS per well overnight at 4°C. Plates were then washed with PBS 0.05% Tween 20™ (ThermoFisher) and blocked in wash buffer with 5% milk (Safeway). Serum complement was heat inactivated at 55°C for 30 minutes and serially diluted in blocking buffer with the least dilution of 1:50. Sera dilutions were then incubated on ELISA plates for 1.5 hours at room temperature. These were then washed and stained with anti-mouse IgG (γ-chain specific)−peroxidase antibody (MilliporeSigma #A3673) diluted 1:10,000 in wash buffer for 1 hour at room temperature. Secondary antibody was then washed and replaced with 100µL of 4µg/mL o-Phenylene diamine in a buffer of 50mM citric acid 100mM dibasic sodium phosphate 0.01% hydrogen peroxide pH 5.0 for 10 minutes before color change reaction was stopped with equal amount of 1M HCl. Absorbance at 490nm was measured using BioTek Synergy HTX Multimode Reader and Gen5 Microplate Reader and Imager Software v3.11 (Agilent). Background absorbance was defined as the lowest absorbance value on a plate and subtracted from each well. Regression curves were fit using Microsoft Excel to determine endpoint titer defined as when absorbance was 0.1. Values < 1 were adjusted to an endpoint titer = 1 for figures. Values below the least dilute serum tested (log(1/50)= 1.7) were set to 1.7 for all statistical analyses.

### Focus reduction neutralization test (FRNT)

Serial dilutions of sera were prepared in DMEM with 2% FBS and incubated with POWV-I for 1 hour at 37°C to allow for antibody binding. Serum treated virus was then used to infect monolayers of Vero cells and overlaid with CMC as described previously. The least dilute serum used to neutralize virus was 1:50. Cells were then fixed with 4% paraformaldehyde at 48 hours post-infection for 30 minutes, blocked and permeabilized in PBS with 2% goat serum (ThermoFisher) and 0.4% Triton™ X-100 (ThermoFisher). Cells were then stained with T077, a monoclonal antibody from a TBEV-infected individual sequenced and characterized by Agudelo *et. al* that binds both POWV-I and -II [42]. We cloned the T077 heavy and light variable chain sequences into pcDNA-3-RhIgG1 and -RhIgK, respectively, for antibody production in Expi293 cells. POWV foci were then visualized by secondary staining with monkey IgG gamma peroxidase-conjugated antibody (Rockland) diluted 1:1,000 in blocking buffer followed by Vector® VIP Substrate Peroxidase (HRP) Kit (Vector Laboratories) after washing. Foci were counted using AID Elispot 7.0 (Autoimmun Diagnostika GMBH) and reciprocal FRNT50s determined by non-linear regression analysis with a variable slope on GraphPad Prism v10.2.2. Values < 1 were adjusted to FRNT50 = 1 for figures. Values extrapolated by GraphPad below the least dilute serum tested (log(1/50)= 1.7) were set to 1.7 for all statistical analyses.

### Area under the curve (AUC) analysis

Areas under the curve of individual mouse weights were measured for curves of percent initial weight from day 0 through 21 using GraphPad Prism v10.2.2.

### Statistical analyses

All appropriate statistical analyses were performed using GraphPad Prism v10.2.2.

### Examination of VLPs by TEM negative stain

Glow discharged, carbon coated Formvar copper grids, 400 mesh, were floated onto 5µl aliquot suspension for 2 minutes. Excess solution was wicked off with filter paper and stained with 1% aqueous uranyl acetate. Stain was removed with filter paper, air dried, and examined on a FEI Tecnai T12 TEM, operated at 80 kV, and digital images were acquired with an AMT Nanosprint12 4k x 3k camera.

## Acknowledgements

Electron microscopy was performed at the Multiscale Microscopy Core, a member of the OHSU University Shared Resource Cores, RRID: SCR_022652. Funding provided by NIH grant R01AI152192 to AJH.

## Supporting information

**S1 Fig.**
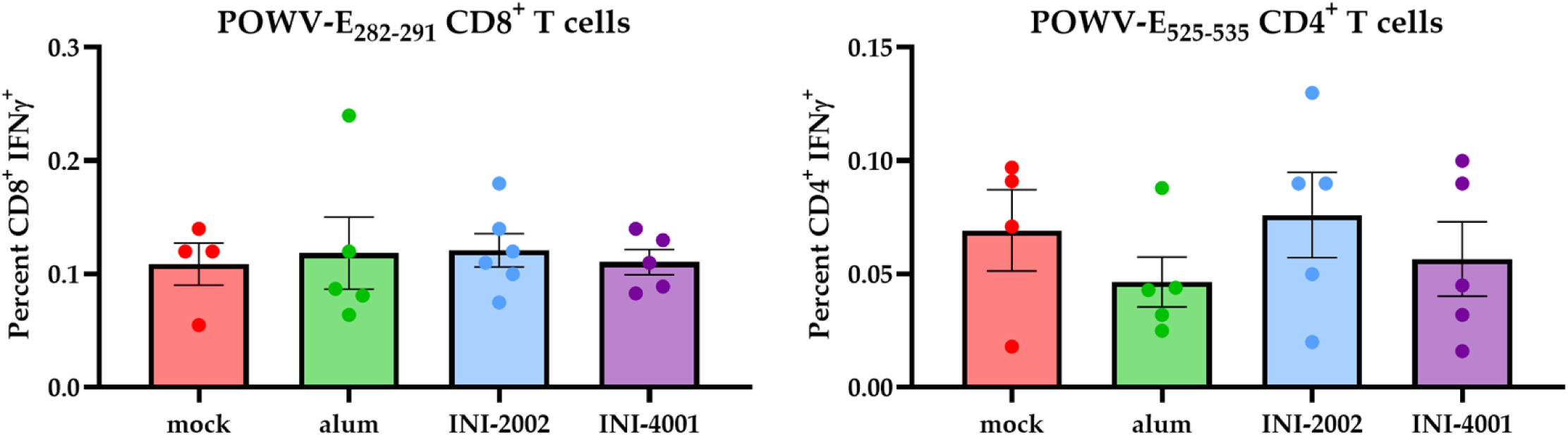
Vaccination does not induce measurable CD8^+^ or CD4^+^ T cell responses. (A-B) Mice prime-boost vaccinated 4 weeks apart *sc* with 10^6^ FFUe of VLP alone or adjuvanted with 300µg alum, 1nmol INI-2002, or 10nmol INI-4001. Mock group injected with PBS vehicle alone. n=4-5. Spleens collected 4 weeks post-boost, and splenocytes incubated with POWV-E_282-291_ (A) or POWV-E_525-535_ (B) for CD8^+^ and CD4^+^ T cells, respectively. Cells were then stained and gated on CD19-CD4+/CD8+ and intracellularly stained for IFNγ. Data are reported as percentage of IFNγ^+^ T cells either within the CD8^+^ (A) or CD4^+^ (B) compartments ± SEM. Data represent one experiment. Statistical significance determined by one-way ANOVA and Šídák’s multiple comparisons test to compare treatments to mock; no statistically significant differences were found.

## Notes

### Competing Interest Statement

DB and JE are employees of and/or shareholders of Inimmune Corp., which holds an exclusive license for the INI-4001 and INI-2002 used in this study. The funders had no role in the design of the study in the collection, analyses, or interpretation of data in the writing of the manuscript or in the decision to publish the results.

